# ATPase SRCAP is a new player in cell division, uncovering molecular aspects of Floating-Harbor syndrome

**DOI:** 10.1101/2020.09.12.294645

**Authors:** Giovanni Messina, Yuri Prozzillo, Francesca Delle Monache, Maria Virginia Santopietro, Maria Teresa Atterrato, Patrizio Dimitri

**Affiliations:** Dipartimento di Biologia e Biotecnologie “Charles Darwin” Sapienza Università di Roma, Roma, Italy; Istituto Pasteur Italia, Fondazione Cenci-Bolognetti

**Keywords:** Cell cycle, Cytokinesis regulators, Midbody, SRCAP, Floating Harbor

## Abstract

Floating-Harbor syndrome (FHS) is a rare genetic disease affecting human development caused by heterozygous truncating mutations in the *Srcap* gene, which encodes the ATPase SRCAP, the core catalytic subunit of the homonymous chromatin-remodeling complex. Using a combined approach, we studied the involvement of SRCAP protein in cell cycle progression in HeLa cells. In addition to the canonical localization in interphase nuclei, both SRCAP and its *Drosophila* orthologue DOMINO-A localized to the mitotic apparatus after nuclear envelope breakdown. Moreover, SRCAP and DOMINO-A depletion impaired mitosis and cytokinesis in human and Drosophila cells, respectively. Importantly, SRCAP interacted with several cytokinesis regulators at telophase, strongly supporting a direct role in cytokinesis, independent of its chromatin remodeling functions. Our results provide clues about previously undetected, evolutionarily conserved roles of SRCAP in ensuring proper mitosis and cytokinesis. We propose that perturbations in cell division contribute to the onset of developmental defects characteristic of FHS.

**Summary:** *Significance statement:* *Srcap* is the causative gene of the rare Floating Harbor syndrome (FHS). It encodes the ATPase SRCAP, the core catalytic subunit of the homonymous multiprotein chromatin-remodeling complex in humans, which promotes the exchange of canonical histone H2A with the H2A.Z variant. According to the current view on SRCAP protein functions, FHS is caused by chromatin remodeling defects. Our findings suggest that, in addition to the established function as epigenetic regulator, SRCAP plays previously undetected and evolutionarily conserved roles in cell division. Hence, we propose that perturbations in cell division produced by SRCAP mutations are important causative factors co-occurring at the onset of FHS.

## Introduction

In the last few decades, a large body of experimental evidence has suggested that mutations in genes encoding a variety of chromatin factors and epigenetic regulators, such as DNA or histone modifying enzymes and members of ATP-dependent chromatin remodeling complexes, are crucial players in human genetic diseases and cancer (Bickmore and van der Maarel, 2003; Bouazoune and Kingston, 2012; Kumar et al., 2016; Masliah-Planchon et al., 2015). Floating-Harbor syndrome (FHS), also known as Pelletier–Leisti syndrome, is a human developmental disorder characterized by delayed bone mineralization and growth deficiency, which are often associated with mental retardation and skeletal and craniofacial abnormalities (Hood et al., 2012; Messina et al., 2016a; Nikkel et al., 2013; White et al., 2010).

*Srcap* (SNF2-Related CBP Activator Protein) is the causative gene of the FHS (MIM number 136140). It maps to chromosome 16p11.2 and encodes the ATPase SRCAP, the core catalytic subunit of the homonymous multiprotein chromatin-remodeling complex in humans. The SRCAP complex is member of the evolutionarily conserved INO80 family of ATP-dependent chromatin remodeling complexes (Bao and Shen, 2011; Clapier and Cairns, 2009) and contains a dozen subunits (Bao and Shen, 2011; Clapier and Cairns, 2009; Havugimana et al., 2012; Messina et al., 2016b; Messina et al., 2017; Messina et al., 2015; Prozzillo et al., 2019; Ruhl et al., 2006). The primary function of the SRCAP complex is to catalyze the incorporation of H2AZ-H2B dimers into nucleosomes (Feng et al., 2018; Wong et al., 2007).

FHS has a dominant inheritance pattern caused by nonsense or frameshift mutations in exons 33 and 34 of *Srcap* gene (Hood et al., 2012; Messina et al., 2016a). These mutations are supposed to produce a C-terminal-truncated SRCAP protein variant missing the AT-hook motifs with DNA-binding activity and are possibly responsible for a dominant negative effect (Hood et al., 2012; Messina et al., 2016a) triggering the onset of FHS. Recently, localization assays in human embryonic cranial neural crest cells showed that overexpressed GFP/Flag-tagged versions of C-terminal-truncated SRCAP are largely excluded from the nucleus but present in the cytoplasm (Greenberg et al., 2019), suggesting that FHS mutations affect the nuclear localization of SRCAP. The SRCAP protein can also function as a transcriptional activator by binding to the cAMP response element-binding protein (CREB)-binding protein (CREBBP or CBP) (Johnston et al., 1999). Moreover, a role of SRCAP in DNA-end resection was proposed recently (Dong et al., 2014).

DOMINO-A (DOM-A), the functional SRCAP orthologue in *Drosophila melanogaster*(Eissenberg et al., 2005; Ruhf et al., 2001), is the main subunit of the *Drosophila* Tip60 (dTip60) chromatin-remodeling complex (Kusch et al., 2004). The dTip60 subunits share high sequence identity and functional conservation with SRCAP and p400/Tip60 human complexes (Clapier and Cairns, 2009).

Overall, SRCAP appears to be a multifaceted protein implicated in several cellular processes, including chromatin regulation, transcription, and DNA repair (Clapier and Cairns, 2009; Dong et al., 2014; Feng et al., 2018; Johnston et al., 1999; Monroy et al., 2001; Wong et al., 2007). Therefore, investigating the wild-type cellular functions of SRCAP may provide clues to the genetic and molecular basis of FHS onset (Messina et al., 2016a).

We combined cell biology, reverse genetics, and biochemical approaches to study the subcellular distribution of the SRCAP protein and assess its involvement in cell cycle progression in HeLa cells. Unexpectedly, we found that SRCAP associates with components of the mitotic apparatus, including centrosomes, the spindle, and midbody, and interacts with a plethora of cytokinesis regulators. Furthermore, its RNAi-mediated depletion perturbs mitosis and cytokinesis. Importantly, SRCAP was found to interact at telophase with a number of cytokinesis regulators and control their midbody recruitment. Similarly, DOM-A localizes to centrosomes and the midbody, and its depletion results in cell division defects in Drosophila S2 cells. Our findings suggest that, in addition to the established role as epigenetic regulator, SRCAP plays previously undetected and evolutionarily conserved roles in cell division. Moreover, our results emphasize a surprising scenario whereby alterations in cell division produced by SRCAP mutations are important causative factors co-occurring at the onset of FHS.

## Results

### SRCAP is recruited to the mitotic apparatus in human cell lines

First, we investigated the subcellular distribution of endogenous SRCAP protein during the cell cycle in HeLa cells using immunofluorescence microscopy (IFM). As shown in Fig. 1, a SRCAP polyclonal antibody (T15, Table S1) decorated the interphase nuclei, as expected, but also revealed a specific pattern at the mitotic apparatus during mitotic progression. Soon after nuclear envelope breakdown, SRCAP immunofluorescence redistributed at the mitotic spindle with enrichment at the poles and centrosomes and later at the central spindle and midbody. These observations were recapitulated in the HuH7 hepatocyte carcinoma-derived cell line (Nakabayashi et al., 1982) and in human MRC5 fibroblast-derived cell line (Fig. S1), in that immunofluorescent signals were observed at centrosomes, spindle and midbody, in line with the results in HeLa cells (Fig. 1). The specificity of the SRCAP antibody was validated by both Western blotting and IFM performed after RNAi-mediated depletion of SRCAP in HeLa cells (Figure S2 and Table S1). Taken together, these results indicate that the observed localization reflects intrinsic properties of the SRCAP protein, with no cell type specificity.

**Fig. 1.**
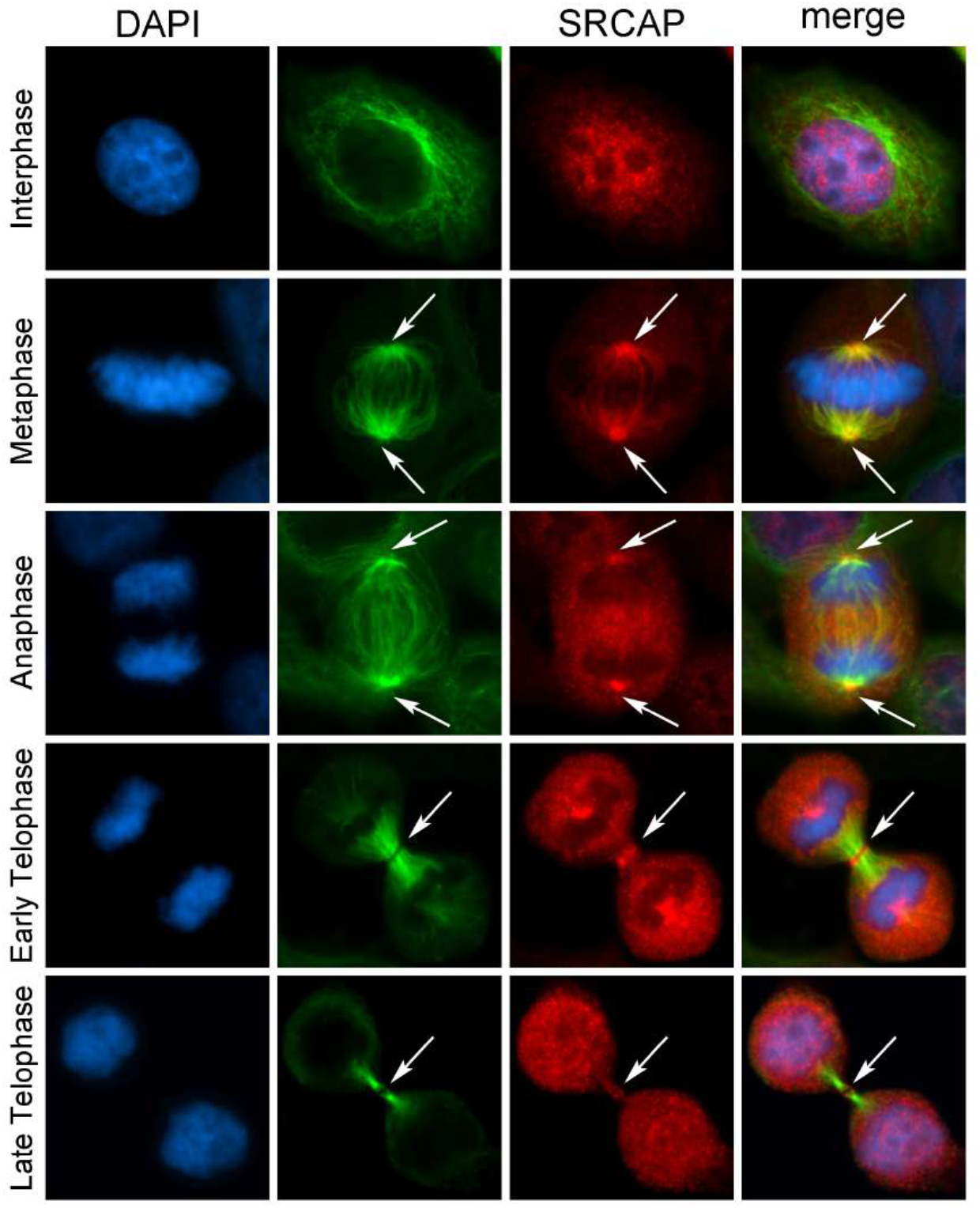
SRCAP localizes to the centrosomes, spindle and midbody in HeLa cells. From left to the right: DAPI (blue), anti-SRCAP (red), anti-α-Tubulin (green) and merge. As expected, the SRCAP staining is clearly present in the interphase nuclei. At metaphase, the SRCAP staining is found on spindle poles and spindle fibers, while in later stage is found on centrosomes and central spindle (anaphase) and midbody (telophase).

The midbody is a tightly packed bridge that forms from the bipolar microtubule array derived from the anaphase central spindle. It serves as a platform for orchestrating cytokinesis by recruiting a large number of factors needed for abscission, the last stage of cell division (Normand and King, 2010). Therefore, we wanted to evaluate the midbody association of SRCAP using both IFM and Western blotting on isolated midbodies (see Materials and Methods). As shown in Fig. 2A, SRCAP immunofluorescence clearly decorated the isolated midbodies. Western blot analysis confirmed the presence of SRCAP protein in extracts from isolated midbodies (Fig. 2B).

**Fig. 2.**
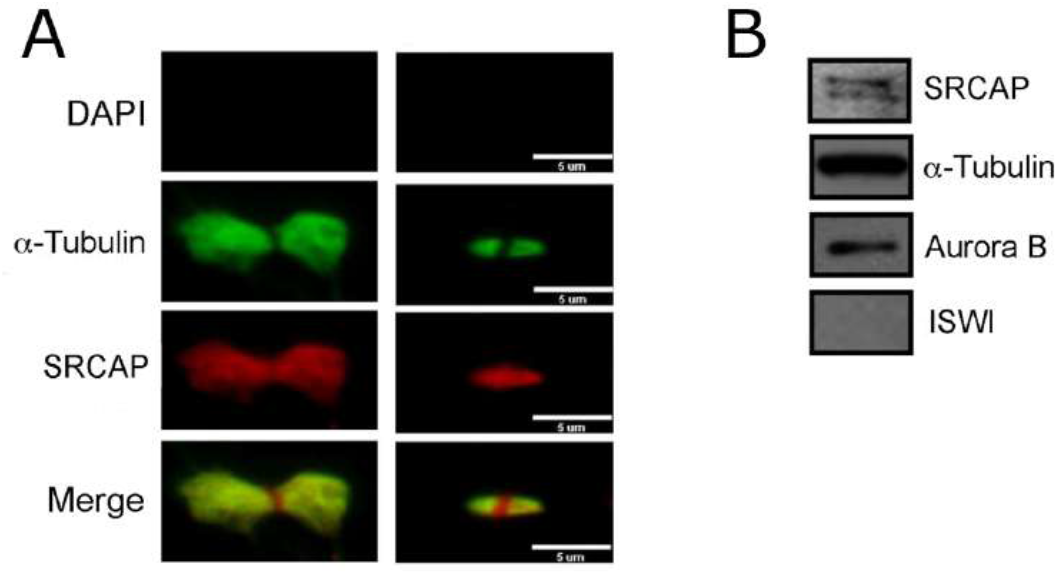
Localization of SRCAP on isolated midbodies using the T-15 SRCAP antibody. Fixed preparations of midbodies isolated from HeLa cells were stained with DAPI (blue), anti-SRCAP (red) and anti-α-Tubulin (green). A) Immunolocalization of SRCAP protein to isolated midbodies. No DAPI staining was detected. SRCAP staining clearly decorated the isolated midbodies and overlaps with that of α-Tubulin. B) Detection of SRCAP by Western blotting on midbody extracts from HeLa cells. Two SRCAP bands were detected of about 350 and 300 KDa. Aurora B was used as a positive control. The ISWI remodeller (negative control) was not detected.

Taken together, these findings show that the subcellular localization of SRCAP is dynamic during cell division and it is recruited not only to interphase nuclei, but also to the centrosomes, spindle, and midbody. Remarkably, SRCAP is the core subunit of the homonymous complex governing H2A.Z deposition into chromatin (Clapier and Cairns, 2009; Feng et al., 2018; Wong et al., 2007), thus, its association with the mitotic apparatus was not obvious.

### SRCAP protein depletion perturbs mitosis and cytokinesis in HeLa cells

Next, we examined the functional significance of SRCAP recruitment to centrosomes, the spindle, and midbody during cell division by investigating the progression of cell division in SRCAP-depleted HeLa cells utilizing RNAi-mediated silencing. In fixed HeLa cell preparations, we categorized and quantified six classes of cell division defects (Fig. 3, A-G and Table 1): multipolar spindles (MS) at pro-metaphase and metaphase (Fig. 3B), chromosome misalignments (CM) and altered spindle morphology (ASM) at metaphase (Fig. 3C), chromatin bridges (CB) at anaphase and telophase (Fig. 3D), long thin intercellular bridges (LIB) at the last stage of telophase (Fig. 3E), and multinucleated cells (MC) (Fig. 3F). Compared to mock-treated control cells, SRCAP RNAi-treated cells exhibited a significant increase in mitosis and cytokinesis defects in all classes except MS. The increase was particularly relevant for CM (57%). Notably, the misaligned chromosomes carry the centromere (Fig. S3), strongly suggesting they were not lost fragments resulting from chromosome breaks. Moreover, a relevant increase of LIB was observed (29%). A LIB is defined as overextended, stretched, intercellular bridge that forms as a consequence of a failure of abscission, the final stage of cytokinesis. Consistently, the intercellular distance at the abscission stage in SRCAP-depleted cells was also increased compared to control cells (Fig. 3G). Defective cytokinesis was also reflected in the appearance of MC (14%), whereas the formation of CB was mild compared to the mock-treated control. Moreover, in agreement with defective mitosis, SRCAP-depleted cells exhibited a significant amount of altered spindles with a shorter and thinner shape (ASM) than those in the control cells (Fig. 3C). Thus, it appears that SRCAP depletion disrupts both mitosis and cytokinesis in HeLa cells, suggesting that the localization observed at centrosomes, the spindle, and midbody reflects its functional roles in cell division.

**Fig. 3.**
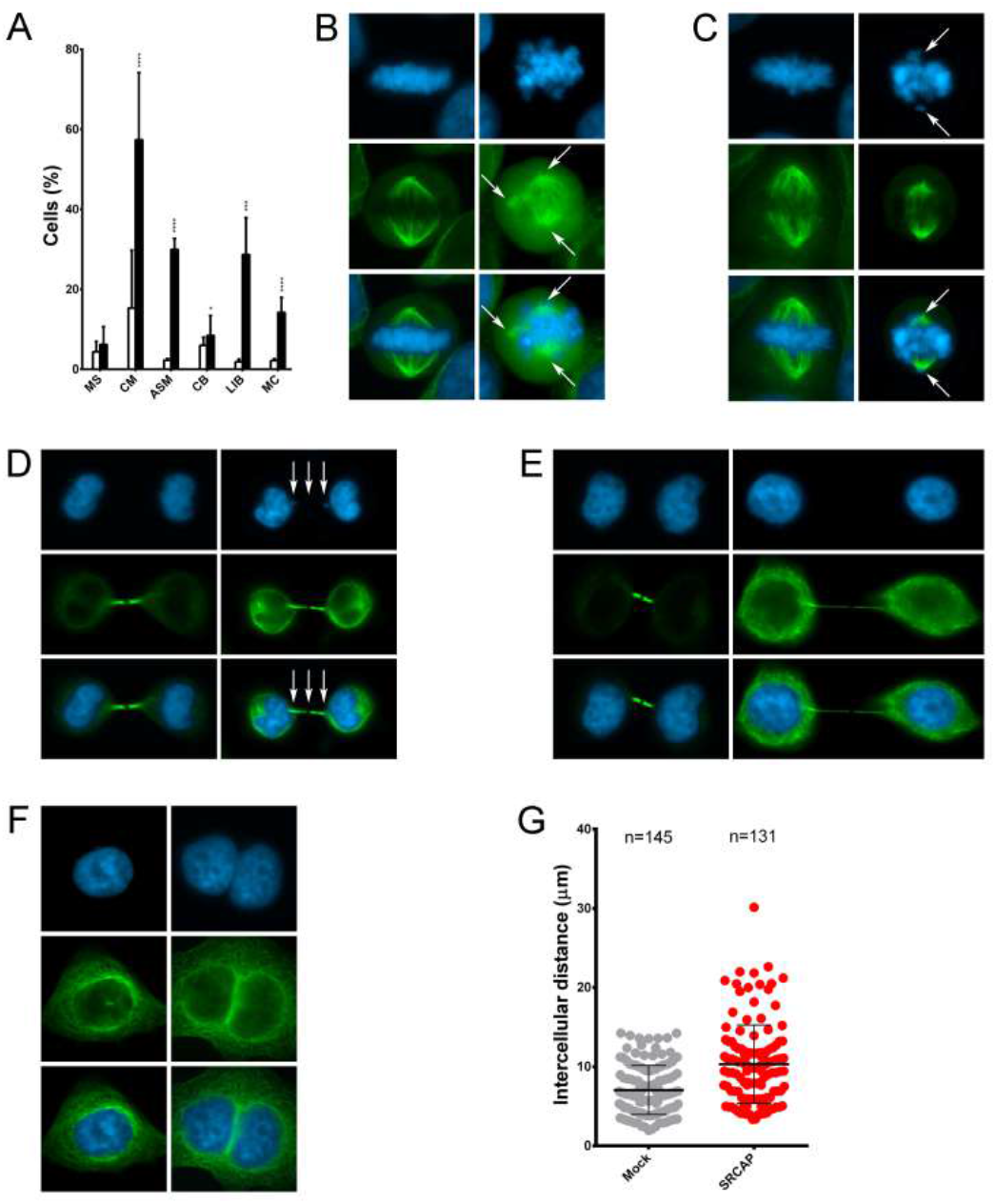
RNAi mediated depletion of SRCAP affects cell division in HeLa cells. RNAi knock-down experiments were performed by transfecting HeLa cells with specific siRNAs (see Materials and Methods). Hela cells were stained with DAPI (blue) and anti-α-Tubulin (green). Left panels (mock), right panels (RNAi). Six classes of defects were categorized: A) Histograms showing the quantitative analysis of cell division defects. B) Multipolar spindles (MS). C) Chromosome misalignments (CM) and altered spindle morphology (ASM). D) Chromatin bridges (CB). E) Long intercellular bridges (LIB); no DAPI-stained trapped chromatin was observed. F) Multinucleated cells (MC). G) Intercellular distance. The quantitative analysis of defects scored in RNAi-treated and control cells (Table 1) is based on the following numbers: at least 100 prometaphases and metaphases for MS, 70 metaphases for CM and ASM, 300 telophases for LIB and CB, 5500 for MS. Three independent experiments were performed.

**Table 1.**
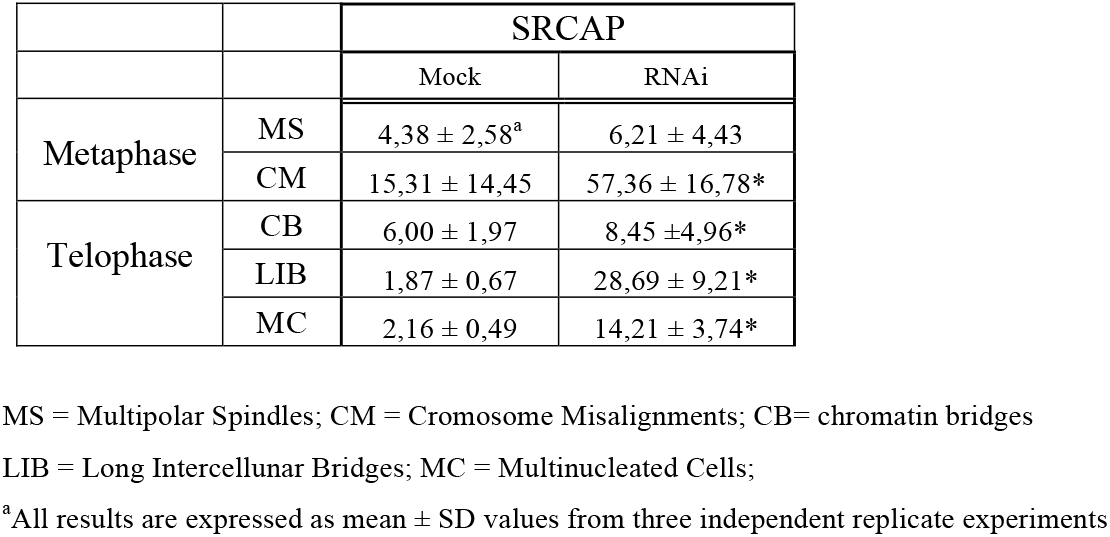
Cell division defects found in SRCAP depleted HeLa cells

|  |  | SRCAP |  |
| --- | --- | --- | --- |
|  |  | Mock | RNAi |
| Metaphase | MS | 4,38 ± 2,58<sup>a</sup> | 6,21 ± 4,43 |
|  | CM | 15,31 ± 14,45 | 57,36 ± 16,78* |
| Telophase | CB | 6,00 ± 1,97 | 8,45 ± 4,96* |
|  | LIB | 1,87 ± 0,67 | 28,69 ± 9,21* |
|  | MC | 2,16 ± 0,49 | 14,21 ± 3,74* |

MS = Multipolar Spindles; CM = Chromosome Misalignments; CB= chromatin bridges
LIB = Long Intercellular Bridges; MC = Multinucleated Cells;
<sup>a</sup>All results are expressed as mean ± SD values from three independent replicate experiments

**Table 2.** Cytokinesis regulators misslocalizations in SRCAP depleted HeLa cells

|  | Mock | SRCAP |
| --- | --- | --- |
| CIT-K | 2,04 ± 2,65<sup>a</sup> | 4,80 ± 0,75 |
| MKLP2 | 3,03 ± 2,90 | 39,84 ± 15,78* |
| Aurora B | 4,81 ± 5,90 | 56,5 ± 26,16* |
| INCENP | 9,28 ± 3,40 | 60,90 ± 11,93* |
| MKLP1 | 0,37 ± 0,57 | 36,78 ± 1,52* |
| PLK1 | 3,86 ± 6,06 | 36,91 ± 27* |
| Cep55 | 3,33 ± 1,15 | 8,27 ± 1,61* |
| Anillin | 11,26 ± 5,66 | 31,38 ± 9,67* |
| Alix | 5,83 ± 1,06 | 29,15 ± 6,48* |
| Spastin | 12,4 ± 3,18 | 29,90 ± 13,43* |

<sup>a</sup>All results are expressed as mean ± SD values from three independent replicate experiments.
\*=P < 0.05, compared with the Mock group

### Microtubule re-polymerization is affected in SRCAP-depleted cells

The finding that SRCAP depletion affects spindle morphology and chromosome alignment at metaphase (Fig. 3, A and C) suggests that this protein regulates microtubule organization and mitotic spindle assembly. To test this hypothesis, we used HeLa cells stably expressing EGFP::α-Tubulin and assessed whether the presence or absence of SRCAP influences microtubule regrowth after cold-induced disassembly. Control (mock) and SRCAP RNAi-depleted HeLa cells (RNAi) were incubated on ice for 1 hour to induce extensive depolymerization (T0). The cells were then allowed to rewarm at 37°C in complete medium for 5 min (T5) to resume microtubule regrowth. As shown in Fig. 4, microtubule repolymerization after 5 minutes of rewarming resulted in clearly aberrant asters with rare, long, and thin MTs in RNA-treated cells (mean±SD, 36.84%± 3.76) compared to the mock-treated cells (mean±SD 5%± 1.92). This result supports a role of the SRCAP protein in microtubule organization and mitotic spindle assembly.

**Fig. 4.**
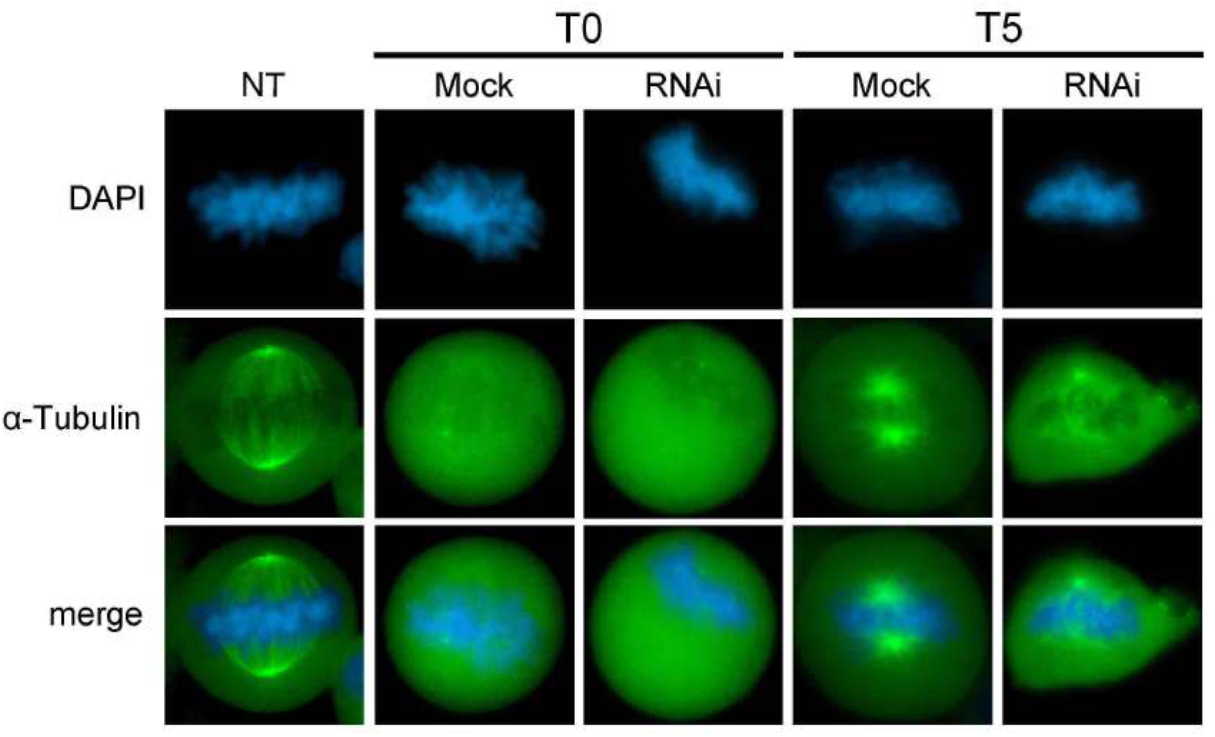
Abnormal microtubules re-polymerization in SRCAP depleted cells. From left to the right: DAPI (blue), α-Tubulin (green) and merge. In control Hela cells microtubule re-polymerization after 5 minutes of rewarming give rise to properly shaped asters, while in SRCAP depleted HeLa cells (RNAi), aster reformation is clearly aberrant with α-Tubulin fluorescence marking only one pole spot, together with sparse and disorganized fibers. The results are based on a total of three experiments; at least 300 cells were scored from both RNAi-treated and control cells.

### SRCAP depletion affects the localization pattern of cytokinesis regulators to the midbody in HeLa cells

Cytokinesis is the last step of cell division and is controlled by a plethora of essential regulators recruited to the midbody during telophase (Barr and Gruneberg, 2007; Bassi et al., 2013; Carlton et al., 2012; Glotzer, 2005; Hu et al., 2012; Normand and King, 2010). The finding that SRCAP-depleted cultures are enriched in LIB and MC (Fig. 3) prompted us to investigate a possible role of SRCAP in cytokinesis. We used IFM to study the recruitment of crucial regulators of cytokinesis to the midbody in HeLa cells proficient or depleted in SRCAP. We focused on CIT-K, MKLP2, Aurora B, INCENP, MKLP1, PLK1, CEP55, Anillin, Alix, and Spastin, nine well-known proteins that localize to the midbody and are required for cytokinesis (Barr and Gruneberg, 2007; Bassi et al., 2013; Carlton et al., 2012; Glotzer, 2005; Hu et al., 2012; Normand and King, 2010). The results of three independent replicates shows that the midbody localization pattern of these factors was impaired in SRCAP depleted HeLa cells, with the exception of CIT-K (Fig. 5). For example, the midbody localization of Aurora B and Anillin was severely affected, while that of PLK1 became more widely distributed.

**Fig. 5.**
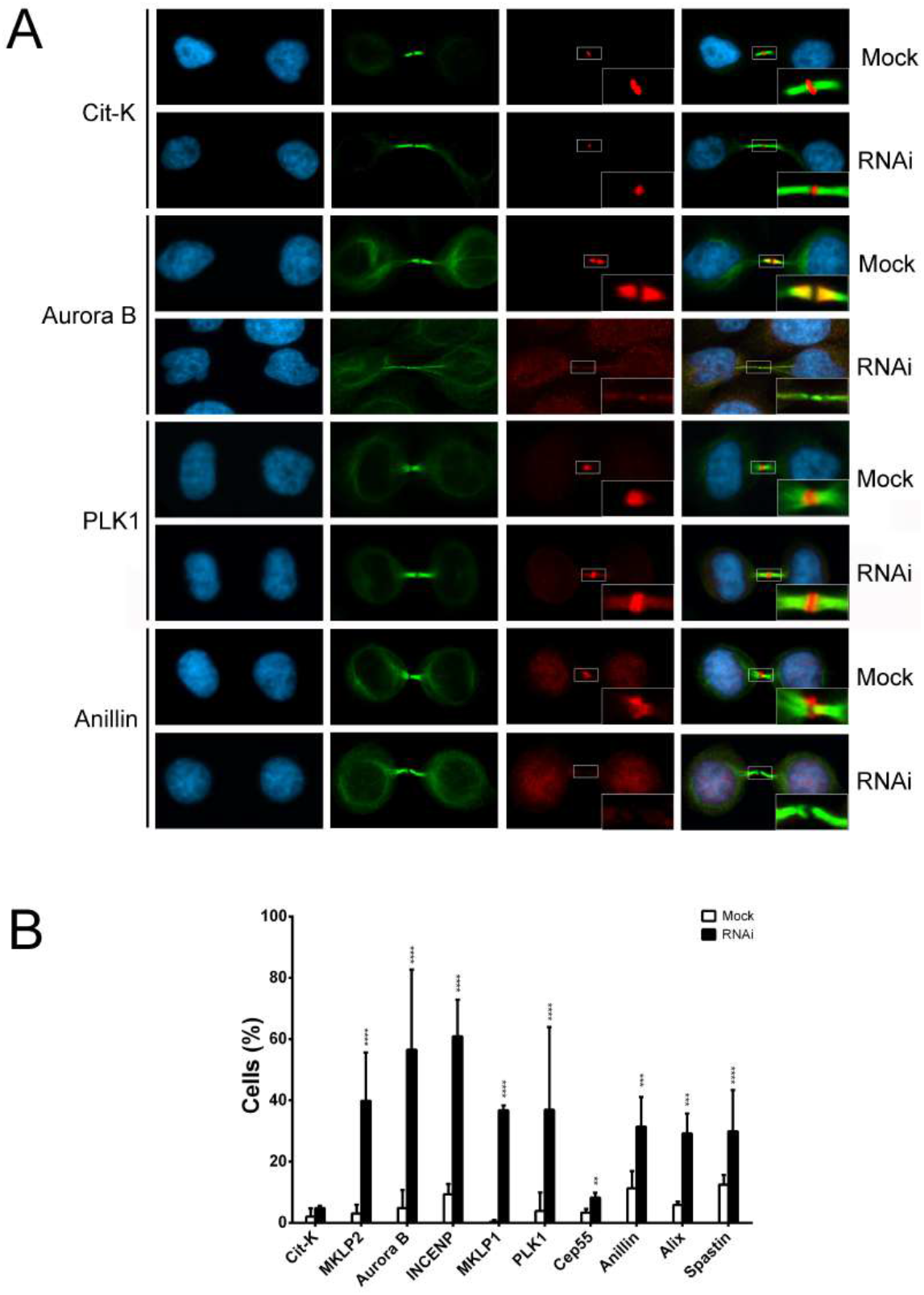
SRCAP depletion affects the midbody localization of cytokinesis regulators. A) Examples of cytokinesis regulators recruitment at midbody in mock and SRCAP depleted cells (RNAi). From left to the right: DAPI (blue), anti-α-Tubulin (green), cytokinesis regulators (red) and merge. B) Histograms showing the quantitative analysis of mislocalizations. Three independent experiments were performed and at least 300 telophases were scored in both RNAi-treated and control cells.

### SRCAP interacts with cytokinesis regulators

The results of the aforementioned experiments (Fig. 5) suggest that the recruitment of a number of cytokinesis regulators to the midbody depends on SRCAP activity. In addition, they are suggestive of possible interactions between SRCAP and the analysed proteins. To test this hypothesis, we carried out co-immunoprecipitation (co-IP) assays on protein extracts from telophase-synchronized HeLa cells. Synchronization was followed by subcellular fractionation assays to recover the cytoplasmic component (S2 fraction) and segregate away the chromatin-associated components (Fig. 6, A and B; Materials and Methods). Fig. 6C shows the comparison between the IP (+SRCAP) and negative control (- SRCAP). SRCAP appears to interact with CIT-K, MKLP2, Aurora B, PLK1, CEP55, Anillin, Alix, and Spastin in telophase. Moreover, we found that SRCAP interacts with α-Tubulin, the main structural component of the midbody. The IP experiments did not reveal any SRCAP interactions with MKLP1 and INCENP (not shown).

**Fig. 6.**
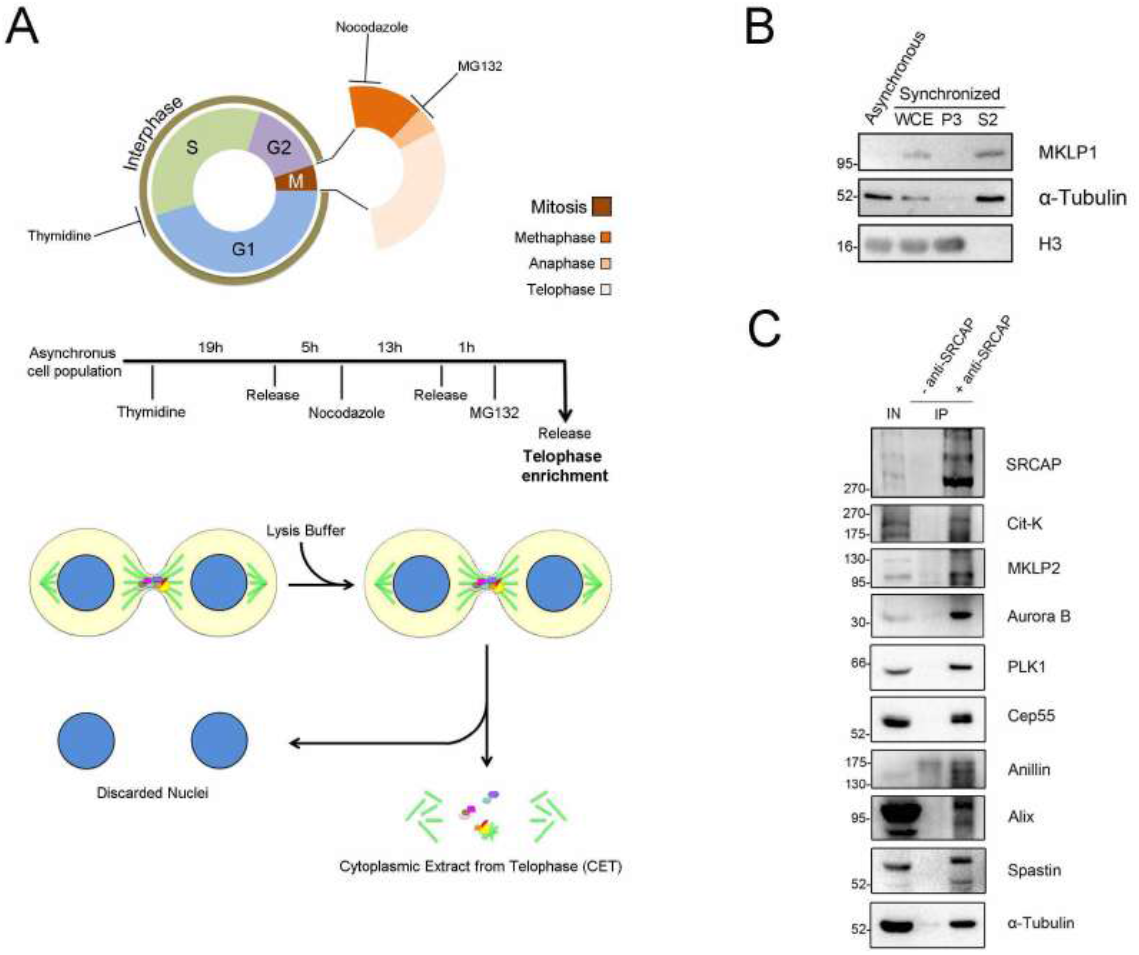
SRCAP interacts with cytokinesis regulators in co-IP assays. We used a SRCAP antibody (Fig. S4 and Table S1) validated by Johnston et al.,1999, since the T-15 antibody did not work in co-IP assays. A) Telophase synchronization in HeLa cells. The scheme summarizes the protocol used for telophase synchronization and subcellular fractionation assay (see Materials and Methods). B) Chromatin fractionation of HeLa cells synchronized in telophase. WCE: whole cell extract. P3: nuclear fraction. S2: cytoplasmatic fraction. H3 and α-Tubulin are markers of nuclear and cytoplasmic fraction, respectively. MKLP1 is expressed in late stages of mitosis (telophase synchronization control). C) Coimmunoprecipitation of protein extracts from telophase synchronized cells (S2 fraction). Three independent IP experiments were performed.

### Aurora B kinase activity inhibition does not affect SRCAP recruitment to the midbody

Our results have shown that SRCAP interacts with Aurora B and contributes to its recruitment to the midbody (Figs. 5 and 6). Aurora B is a key regulator of cell division, and its kinase activity promotes the recruitment of several motor and microtubule-associated proteins in the last stages of cell division and controls abscission when the midbody must is severed (McKenzie et al., 2016; Minoshima et al., 2003; Yan et al., 2010). To gain clues about the factors controlling SRCAP recruitment to the midbody, we treated HeLa cells with ZM447439, a specific selective inhibitor of Aurora B kinase activity (Girdler et al., 2006). The ZM447439 treatment impaired the localization of MKLP1 at the midbody (Fig. 7 and Table 3), a well-known target of Aurora B phosphorylation during the final stage of cell division (Guse et al., 2005). In contrast, the localization of SRCAP was not significantly affected. This result indicates that, although Aurora B interacts with SRCAP by IP, its kinase activity is not responsible for the recruitment of SRCAP to the midbody.

**Fig. 7.**
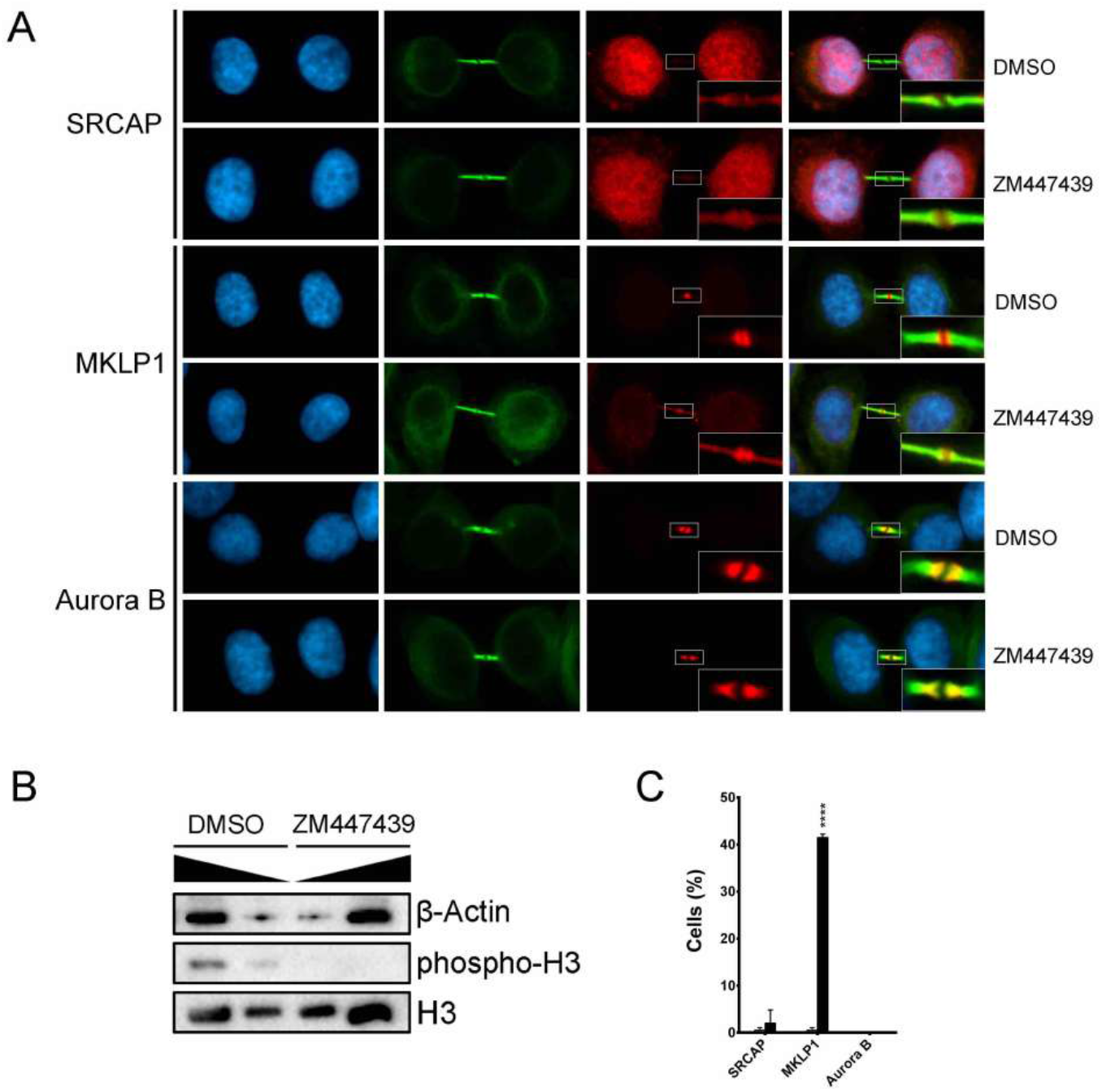
Inhibition of Aurora B kinase activity with ZM447439 does not affect the localization of SRCAP to the midbody. A) From left to the right: DAPI (blue), anti-α-Tubulin (green) and SRCAP, MKLP1 or AURORA-B (red) and merge. Anti-SRCAP immunostaining of ZM447439-treated HeLa cells versus DMSO-treated control HeLa cells (upper panel); anti-MKLP1 immunostaining, positive control (middle panel); anti-Aurora B immunostaining, negative control (lower panel). B) To test the efficacy of ZM447439 inhibition, phosphorylation of the histone H3, a target of Aurora B kinase, during mitosis was evaluated. C) Graph showing mis-localization of MKLP1 to the midbody in ZM447439-treated HeLa cells; no effect on SRCAP and Aurora B localizations was found. At least of 300 telophases were scored in three independent experiments for both treated cells and control HeLa cells.

**Table 3.** Treatment with ZM447439 inhibitor

|  | DMSO | ZM447439 |
| --- | --- | --- |
| SRCAP | 0,50 ± 0,58<sup>a</sup> | 2,00 ± 2,83 |
| MKLP1 | 0,50 ± 0,58 | 41,50 ± 0,71* |
| Aurora B | 0 | 0 |

<sup>a</sup>All results are expressed as mean ± SD values from at least three independent replicated experiments.
\*=P < 0.05, compared with the Mock group.

### Localization and RNAi-mediated depletion of DOM-A in Drosophila S2 cells

Lastly, we investigated whether the association of SRCAP with the mitotic apparatus and defects in cell division observed after its depletion are unique to human cells or are evolutionarily conserved. First, we used IFM to study the localization of DOM-A, the *Drosophila* ortholog of human SRCAP, in *Drosophila melanogaster* S2 cells. In addition to the interphase nuclei, a DOM-A antibody (Ruhf et al., 2001) decorated centrosomes and the midbody (Fig. 8). Next, we examined the phenotypes of S2 cells after RNAi against DOM-A. The RNAi efficiency was tested by immunofluorescent assays and sqRT-PCR (Fig. S5), since the DOM-A antibody did not work properly for Western blotting under our conditions. Depletion of DOM-A resulted in mitotic phenotypes comparable to those observed in SRCAP-depleted HeLa cells (Fig. 9 and Table 4). Five categories of significant defects were observed: MS (46%), CM (21%), CB (4%), LIB (19%), and MC (12%). Importantly, these defects are consistent with the localization of DOM-A to centrosomes and the midbody in S2 cells.

**Fig. 8.**
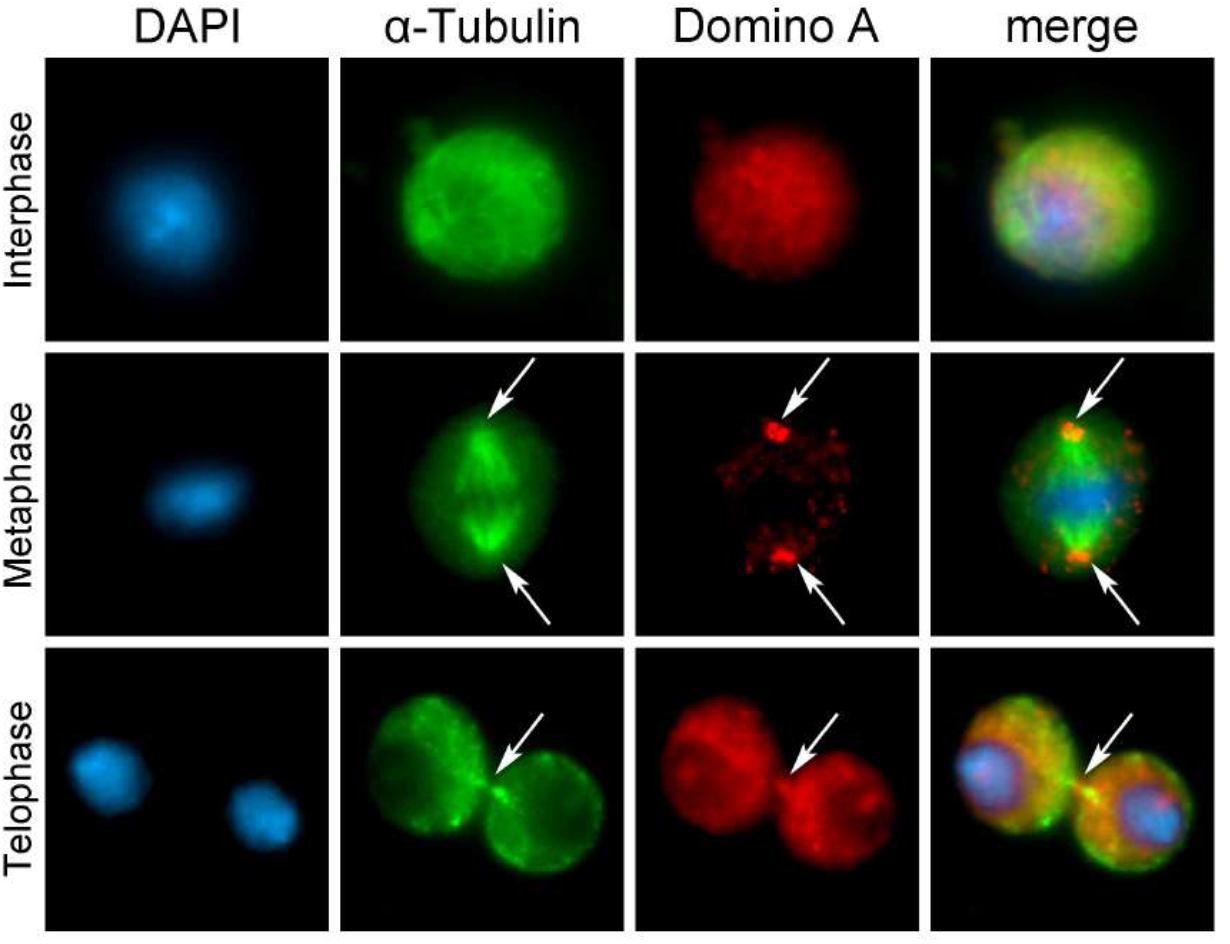
DOM-A localizes to centrosomes and midbody in S2 cells. From left to the right: DAPI (blue), anti-α-Tubulin (green), anti-DOM-A (red) and merge. In addition to interphase nuclei, the anti-DOM-A staining was found on centrosomes (metaphase) and midbody (telophase) pointed by an arrow.

**Fig. 9.**
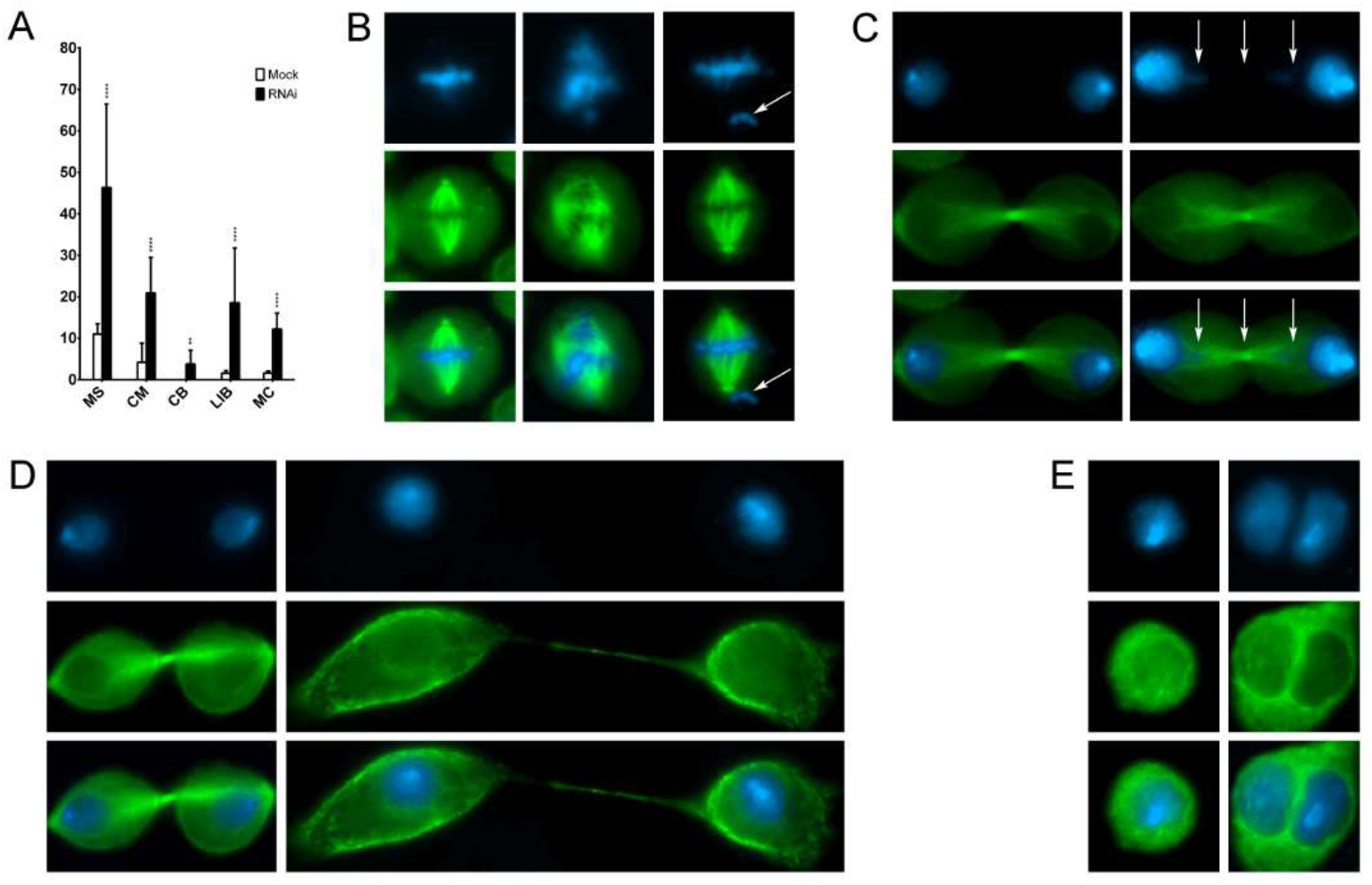
DOM-A depletion in S2 cells affects mitosis and cytokinesis. Examples of mitotic defects found in DOM-A depleted S2 cells and their and quantification. RNAi mediated depletion of DOM-A is described in Material and Methods. DAPI staining is shown in blue, α-Tubulin in green. Square B, from left to right: mock, RNAi, RNAi. Squares C, D and E: left panel mock, right panel RNAi. Five classes of defects were considered: A) Quantitative analysis defects; B) multipolar spindle (MS) and chromosome misalignments (CM); C) chromatin bridges (CB); D) long intercellular bridges (LIB) E) multinucleated cells (MC). The quantitative analysis of defects scored in RNAi-treated and mock treated cells is based on about 300 metaphases, telophases or interphases, scored in at least three independent experiments (see Table 1).

**Table 4.** Quantitative analysis of defects found in DOM-A depleted S2 cells

|  |  | DomA |  |
| --- | --- | --- | --- |
|  |  | Mock | RNAi |
| Metaphase | MS | 11,01 ± 2,48<sup>a</sup> | 46,33 ± 20,14* |
|  | CM | 4,22 ± 4,58 | 20,9 ± 8,6* |
| Telophase | CB | 0 | 3,81 ± 0,50* |
|  | LIB | 1,57 ± 0,6 | 18,54 ± 13* |
|  | MC | 1,16 ± 0,5 | 12,21 ± 3,84* |

<sup>a</sup>All results are expressed as mean ± SD values from three independent replicated experiments.
\*=P < 0.05, compared with the Mock group.

Next, stained DOM-A-depleted S2 cells with an antibody against Spd2, a Drosophila centriole protein used as a centrosome marker (Giansanti et al., 2008). We observed a high percentage of metaphase with multiple centrosomes exhibiting MT-nucleation ability, which gives rise to

MS (Fig. S6). Multiple centrosomes may arise from aberrant centriole disengagement/amplification, which in turn leads to the formation of MS and chromosome mis-segregation (Marteil et al., 2018). Alternatively, abnormal numbers of centrosomes can be a consequence of cytokinesis failure that results in the formation of MC.

## Discussion

Sparse experimental evidence in the literature provides clues for specific roles of chromatin remodeling factors in cell division that are distinct from their functions in chromatin regulation (Corona et al., 1999; Ducat et al., 2008; Gartner et al., 2003; Gentili et al., 2015; Mo et al., 2016; Sigala et al., 2005; Sillibourne et al., 2007; Yokoyama et al., 2009; Zhang et al., 2012).

Here, we demonstrated that ATPase SRCAP, the core subunit of the SRCAP complex, relocates to centrosomes, the spindle, and midbody during cell cycle progression, with its depletion yielding an array of aberrant outcomes of mitosis and cytokinesis (Figs. 1 and 3). Similarly, DOM-A is found at centrosomes and the midbody in *Drosophila* S2 cells, and its depletion affects both mitosis and cytokinesis (Figs. 8 and 9). Notably, defects in cell division resembling those found after SRCAP and DOM-A depletion occur as a consequence of the loss of crucial regulators of spindle and/or midbody organization or function (Barr and Gruneberg, 2007; Bassi et al., 2013; Carlton et al., 2012; Glotzer, 2005; Hu et al., 2012; Normand and King, 2010).

### How SRCAP and DOM-A depletions affect mitosis and cytokinesis?

At least two alternative hypotheses can be considered to explain the defects found after SRCAP and DOM-A depletion. First, a lack SRCAP or DOM-A may give rise to aberrant chromatin changes that alter the expression of genes involved in cell division and/or to perturbations in kinetochore and spindle organization and function. According to this hypothesis, the cell division defects caused by SRCAP and DOM-A depletion are indirect, and the recruitment of SRCAP or DOM-A to the mitotic apparatus only reflects a passive accumulation of disposable factors. Second, SRCAP and DOM-A proteins are essential components of the mitotic apparatus and regulate cell division independent of their functions in chromatin regulation. If this is true, SRCAP and DOM-A depletion directly affects mitosis and cytokinesis.

Several lines of evidence favor the last hypothesis. First, the recruitment of SRCAP and DOM-A to the mitotic apparatus, together with the disruption of specific steps of mitosis and cytokinesis caused by their depletion in humans and *D. melanogaster*, respectively (organisms separated by about 800 myr), are evolutionarily conserved phenomena. Second, the observed defects do not appear to simply be a chaotic disruption of cell division as one would expect by simultaneous up- or down-regulation of a group of genes encoding regulators of cell division due to epigenetic alterations. In contrast, SRCAP and DOM-A depletion yields to an array of aberrant mitotic outcomes falling into specific categories of alterations in mitosis and cytokinesis. Notably, such alterations are consistent with the localization of SRCAP and DOM-A proteins to the mitotic apparatus and also occur after the loss of key regulators of mitosis and cytokinesis (Barr and Gruneberg, 2007; Bassi et al., 2013; Carlton et al., 2012; Glotzer, 2005; Hu et al., 2012; Normand and King, 2010). Finally, and most importantly, a direct role of SRCAP in the final stage of cell division, independent from its chromatin functions, is primarily supported by the recruitment of MKLP2, Aurora B, PLK1, CEP55, Anillin, Alix, and Spastin to the midbody being affected by SRCAP depletion (Fig. 5) and the same proteins interacting with SRCAP in co-IP assays of chromatin-free protein extracts from the cytoplasmic fraction of telophase-synchronized HeLa cells (Fig. 6). Notably, the interaction between SRCAP and Anillin has also been highlighted in a recent study on the midbody interactome (Capalbo et al, 2019). Among the identified SRCAP interactors, Cit-K was not delocalized after SRCAP depletion (Fig. 5), suggesting that Cit-K may acts in telophase upstream of SRCAP.

All of the proteins identified here as SRCAP interactors in telophase (Fig. 6) are essential for successful cell division in different organisms, as their depletion results in aberrant cytokinesis. Cit-K is the main abscission regulator capable of physically and functionally interacting with the actin-binding protein Anillin, a crucial component of the contractile ring and midbody (Giansanti et al., 1999; Normand and King, 2010; Piekny and Glotzer, 2008); MKLP2 is a motor kinesin that binds microtubules and is required for Aurora B recruitment at the central spindle (Gruneberg et al., 2004; Normand and King, 2010). CEP55 recruits Alix at the midbody. Notably, in the absence of CEP55, a series of late-acting abscission factors fail to concentrate at the midbody, including Aurora B, MKLP2, Plk1, PRC1, and ECT2 (Zhao et al., 2006), and the ESCRT machinery. Spastin is a key player in microtubule severing, ensuring the final cut at the midbody, whereas α-Tubulin is a major component of spindle and midbody microtubules.

Although we cannot completely rule out a contribution of chromatin perturbations in cell division defects, our results suggest that SRCAP and DOM-A, similarly to other chromatin remodelers (Corona et al., 1999; Ducat et al., 2008; Gartner et al., 2003; Gentili et al., 2015; Mo et al., 2016; Sigala et al., 2005; Sillibourne et al., 2007; Yokoyama et al., 2009; Zhang et al., 2012), are multifaceted proteins that, in addition to their canonical functions in interphase, may play conserved roles in mitosis and cytokinesis.

In particular, cell division alterations (Fig 3), spindle reformation defects (Fig. 4), and mislocalization of cytokinesis regulators at the midbody (Fig. 5) found in SRCAP-depleted cells, together with specific interactions detected in telophase-synchronized cells (Fig. 6) provides evidence that SRCAP participates in two different steps of cell division: i) it may ensure proper chromosome segregation, regulating microtubule organization and mitotic spindle assembly, and ii) it may be required for midbody function during abscission, participating in the recruitment of cytokinesis regulators to the midbody, ensuring the final cut that is essential for proper abscission. In this context, crosstalk between SRCAP and cytokinesis regulators may occur to control midbody architecture and function similar to the examples found in the literature (McKenzie et al., 2016).

Several ATPases, such as Katanin, Cdc48/p97, ISWI, VPS4, and Spastin, interact with microtubules and play direct roles in mitosis and cytokinesis (Cao et al., 2003; Joly et al., 2016; Yang et al., 2008). The ESCRT-III complex protein CHMP1B recruits Spastin to the midbody (Yang et al., 2008). Intriguingly, depletion of Spastin results in cytokinesis failure phenotypes similar to those found in SRCAP-depleted cells (Connell et al., 2009). Thus, the ATPase activity of SRCAP may be required for its function in mitosis and cytokinesis. In particular, during the final stage of cell division, SRCAP may act as a platform to recruit/stabilize key cytokinesis/abscission regulators to the midbody to ensure the final cut. The precise mechanism underlying the roles of SRCAP in cell division will be an important topic for further analysis.

In conclusion, our results reveal the existence of a previously undetected and evolutionarily conserved phenomenon, whereby SRCAP is recruited to the mitotic apparatus during cell cycle progression in human cell lines (Figs. 1 and S2) and has functional relevance in cell division preventing genetic instability. Therefore, we propose that mitosis and cytokinesis failure are important causative factors co-occurring at the onset of developmental defects characteristic of FHS. It is well known that defective mitosis or cytokinesis can cause chromosomal instability leading to genetically unstable states, hence activating tumorigenic transformation (Ben-David and Amon, 2020; Lens and Medema, 2019). A first case of a tumor associated with FHS has indeed been reported in 2009 (Nelson et al., 2009). Thus, it might be possible that FHS patients also exhibit some predisposition for tumor development. If this was true, then FHS patients should be subjected to clinical trials for cancer prevention.

## Materials and Methods

### Cell Cultures, transfections and RNAi treatments

HeLa cells, purchased from ATTC company, were cultured in 6-well plates in Dulbecco’s modified Eagle’s medium (DMEM) supplemented with 10% FBS (Corning) and a penicillin/streptomycin solution (Gibco, 15140122). RNAi mediated depletion of SRCAP was performed by double transfection (24h + 48h after seeding) with specific siRNA mix targeting SRCAP transcripts (sc-93293, Santa Cruz), using Lipofectamine RNAiMAX transfection reagent (Thermo Scientific), according to the manufacturer’s protocol; 24h after the second transfection cells were harvested for cytological and immunoblotting analysis. Control samples were treated in the same without addition of siRNA.

*Drosophila melanogaster* S2 cells were cultured at 25°C in Schneider’s *Drosophila* Medium (Biowest). RNAi treatments were carried out according to Somma et al. (2002). To perform DOM-A RNAi, each culture was inoculated with 15 μg of specific siRNA. Control samples were treated in the same way without addition of dsRNA. Both dsRNA-treated and control cells were grown for 96h at 25°C and then processed for either immunofluorescence or blotting analysis. To prepare dsRNA, individual gene sequences were amplified by PCR from genomic DNA obtained from first-instar larvae of a wild type *D. melanogaster* strain. The primers used in the PCR reactions were 48 nt base long and all contained a 5’ T7 RNA polymerase binding site (5’-GAATTAATACGACTCACTATAGGGAGAC-3’) joined to a DOM-A specific sequence. The sense and antisense gene-specific DOM-A primers were as follows: for-TCTGGTGCTCAGATCGTGTC; rev-GTTGTCTGCAGCACCTTCAA.

### Cytology and immunostaining

Cytology and immunostaining of HeLa cells were performed according to Messina et al. (2017). Cytology and immunostaining of *Drosophila melanogaster* S2 cells were performed according to Somma et al. (2002).

### Microtubules re-polymerization assays

HeLa Kyoto EGFP-α-Tubulin/H2B-mCherry cell line was purchased from CLC (EMBL, Germany). Cell cultures and transfection were performed according to the previous section; 24h after last transfection, cells were assayed for microtubules re-polymerization. Control (Mock) and SRCAP RNAi-depleted cells (RNAi) were incubated 1h in ice (T0) and then supplemented with complete medium for 5’ (T5) to resume microtubules polymerization at 37°C. Asters length was evaluated for analysis using the ImageJ software.

### Western blotting and immunoprecipitation

Western blotting was performed according to Messina et al. (2017). Primary antibodies and HRP-conjugated secondary antibodies were described in Table S1 and Table S2, respectively. SRCAP protein immunoprecipitation was performed according to Messina et al. (2017), using rabbit anti-SRCAP antibody by Kerafast (Johnston et al., 1999). Cytosolic fraction (2 mg/ml) from subcellular fractionation assay (see the next paragraph) was used as input (IN). As negative control, no antibody was added to a same amount of IN and beads (Santa Cruz Biotechnology).

### Midbody isolation

The midbody association of SRCAP was also evaluated on isolated midbodies. Midbody isolation was performed according to McKenzie et al. (2016). IFM and Western blotting were performed as described in the above paragraphs.

### Inhibition of Aurora B kinase activity

HeLa cells were treated for 35’ with ZM447439 Aurora B inhibitor at a final concentration of 5 μM and then fixed and stained. An equal volume of DMSO was used as negative control.

### Cell cycle synchronization and subcellular fractionation assay

For immunoprecipitation experiments, HeLa cells were synchronized in telophase using thymidine/nocodazole blocks. Cells were treated with 2 mM thymidine (Sigma, T9250) for 19h, released from G1/S block in fresh media for 5h, incubated with 40 nM nocodazole (Sigma, M1403) for 13h and harvested by mitotic shake-off. Mitotic cells were washed three times with PBS and released in fresh medium for 70’ before harvesting and freezing in liquid nitrogen. Telophase cells (2 x 10^7^) were prepared by resuspending in 1 mL of Buffer A for subcellular fractionation according to Messina et al. (2016b).

### Microscope image acquisition

Both human and *Drosophila melanogaster* slides were analyzed using a computer-controlled Nikon Eclipse 50i epifluorescence microscope equipped with UV-1A EX 365/10 DM 400 BA 400, FITC EX 465-495 DM 505 BA 515-555 and TRITC EX 540/25 DM 565 BA 605/55 filters using Plan Achromat Microscope Objective 100XA/1.25 Oil OFN22 WD 0.2 objective and QImaging QICAM Fast 1394 Digital Camera, 12-bit, Mono. Images were imported into ImageJ software (http://rsbweb.nih.gov/ij/) and adjusted for brightness and contrast uniformly across entire fields where appropriate. The figures were constructed in Adobe Photoshop. Fluorescence intensity was assessed using the ImageJ software.

### Statistical analysis

Data analyses were performed using the GraphPad Prism software (GraphPad Software, Inc., La Jolla, CA, USA). All results are expressed as mean **±**SD values from three independent replicate experiments. P value lower than 0.05 (*P < 0.05) was considered to be statistically significant, using two-tailed Fisher’s exact test.

## Online supplemental material

Fig. S1 shows SRCAP localization in HuH7 and MRC5 cell lines. Fig. S2 shows the efficiency of RNAi mediated depletion of SRCAP. Fig. S3 shows the IF staining of SRCAP depleted metaphases with CREST antibody. Fig. S4 shows the validation of Kerafast anti-SRCAP antibody used in co-IP assays. Fig. 5 shows the efficiency of RNAi mediated depletion of DOM-A protein in Drosophila S2 cells. Fig. S6 shows anti-Spd2 staining of DOM-A depleted and control cells.

## Acknowledgments

The authors thank Patrizia Lavia for her comments on the manuscript. The authors are grateful to Tatsuya Hirano for gift of ISWI and phospho-H3 antibodies and to Thomas Mayer for MKLP2 antibody. This work was supported by grants of Istituto Pasteur Italia - Fondazione Cenci Bolognetti and PRIN 2017 (project number 2017FNZRN3).

## Author contributions

PD and GM conceptualized and supervised the study; PD wrote the original draft; GM, YP, FDM, MVS and MTA performed the experiments.

## SUPPLEMENTAL FIGURES

**Figure S1.**
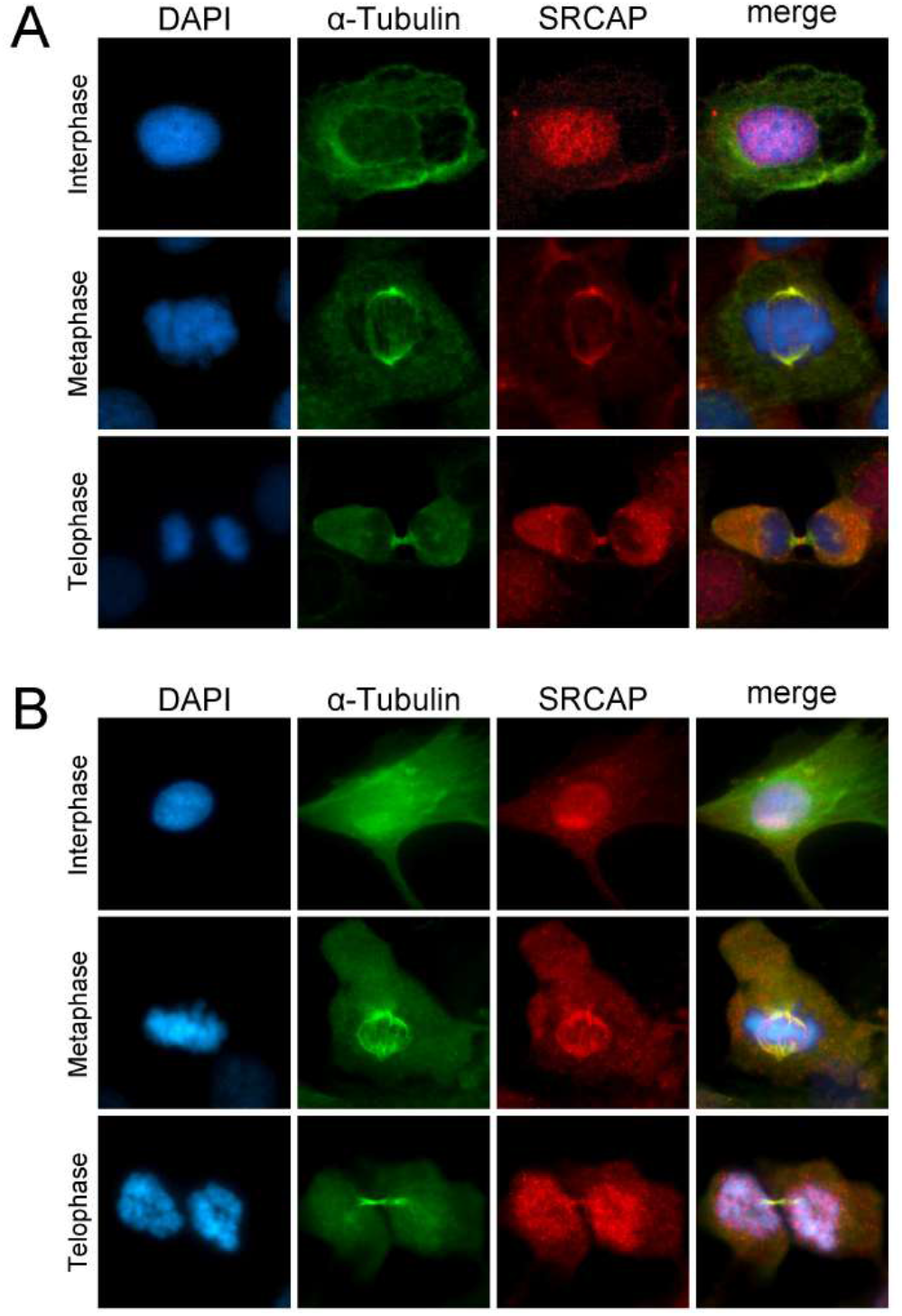
SRCAP localizes at centrosome, spindle and midbody in HuH7 and MRC5 cell lines. Fixed HuH7 (A) and MRC5 (B) cells stained with DAPI (blue), anti-SRCAP (red) and anti-α- Tubulin (green). At interphase, the SRCAP staining decorates the nuclei, while during metaphase and telophase is found at spindle and midbody, respectively.

**Figure S2.**
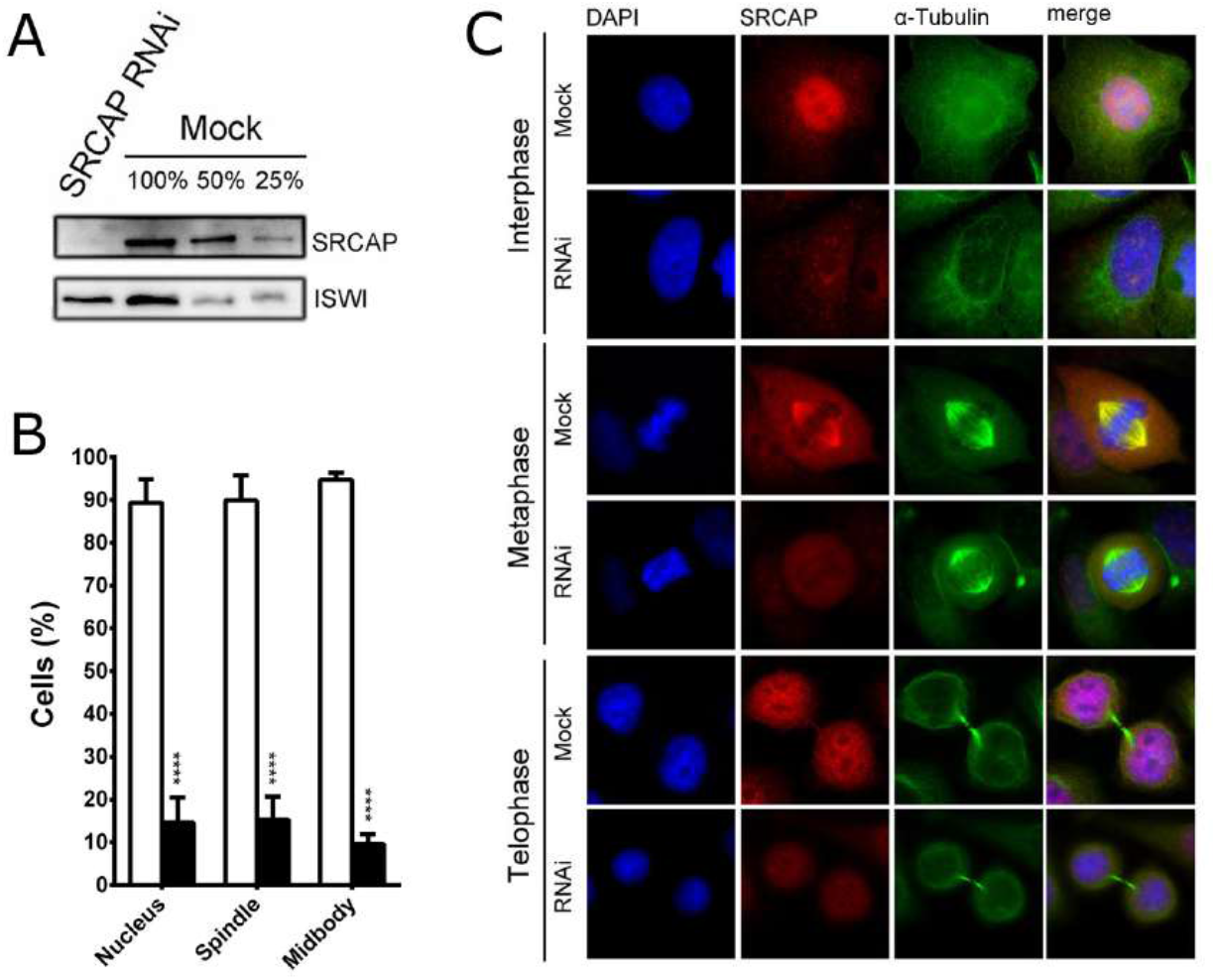
Validation of SRCAP antibody (T15) by IFM and Western blotting. RNAi mediated depletion of SRCAP is described in Materials and Methods. The decrease of SRCAP protein levels after RNAi was measured by IFM and Western blotting, compared to control samples transfected with neutral siRNAs. A) WB of HeLa whole cell extracts after RNAi-depletion of SRCAP. A SRCAP band detected in mock-treated cells was absent from RNAi-treated cells. The ISWI remodeler (negative control) was not significantly affected. B, C). The SRCAP staining shows about 70% decreased in RNAi-treated HeLa cells compared to the mock. Fluorescence intensity was assessed using the ImageJ software.

**Figure S3.**
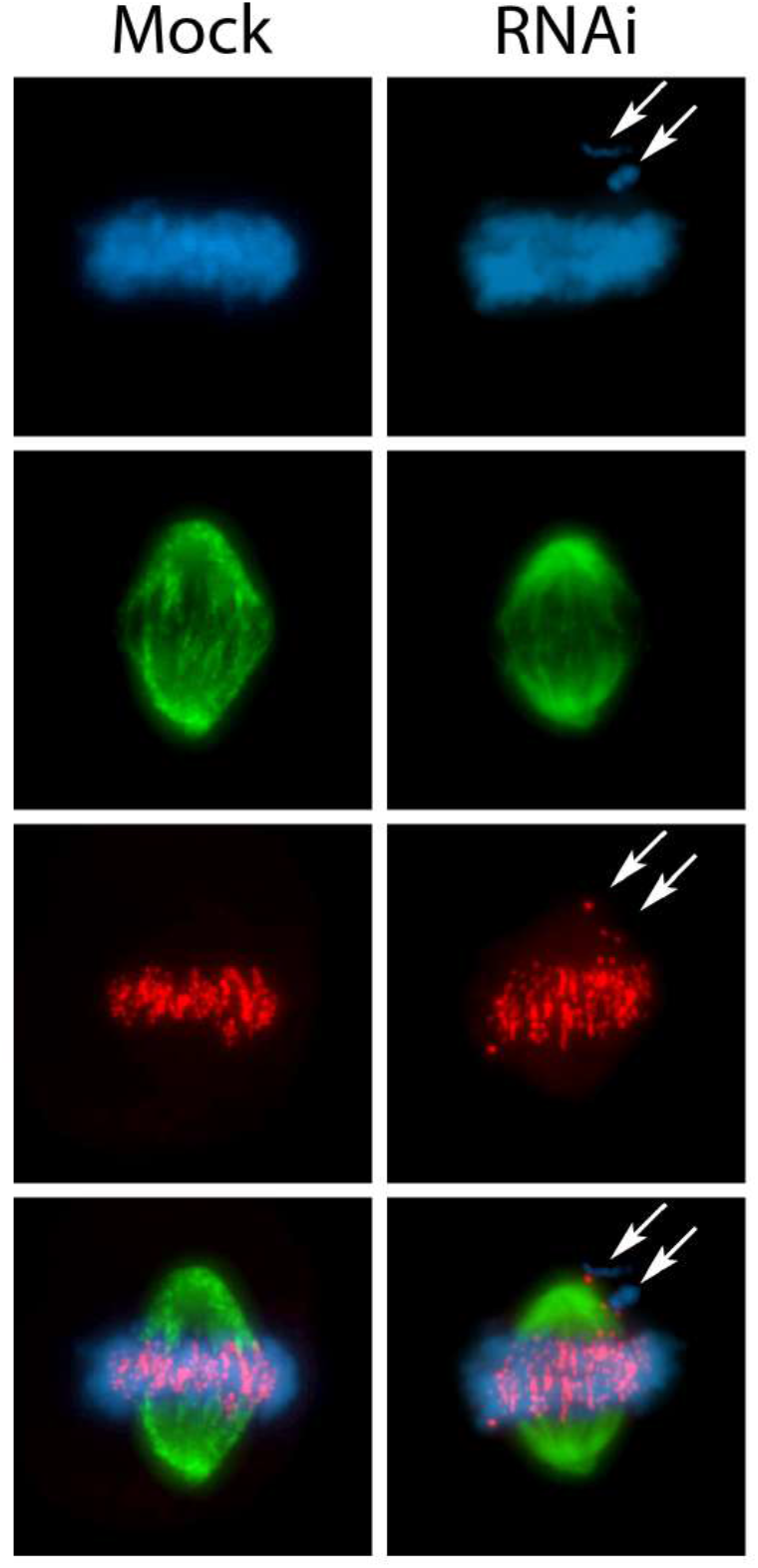
IF assays with CREST antibodies on mock and SRCAP depleted metaphases. The analysis of 300 metaphases scored in three independent experiments showed that all misaligned chromosomes detected carry the centromere.

**Figure S4.**
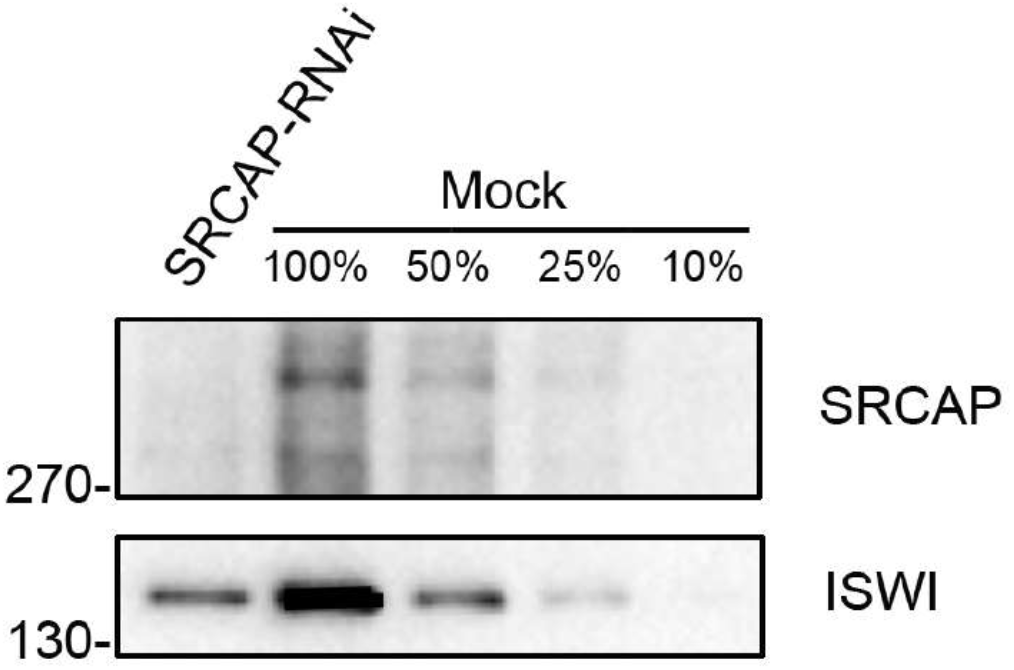
Validation of Kerafast anti-SRCAP antibody used for IP assays. The specificity of Kerafast SRCAP antibody (13) was tested by Western blotting on protein extracts from HeLa cells transfected with specific SRCAP siRNAs (see Materials and Methods), compared to control samples transfected with neutral siRNAs. The SRCAP bands are not found in SRCAP depleted HeLa cells (SRCAP-RNAi). The ISWI remodeler (negative control) was not significantly affected.

**Figure S5.**
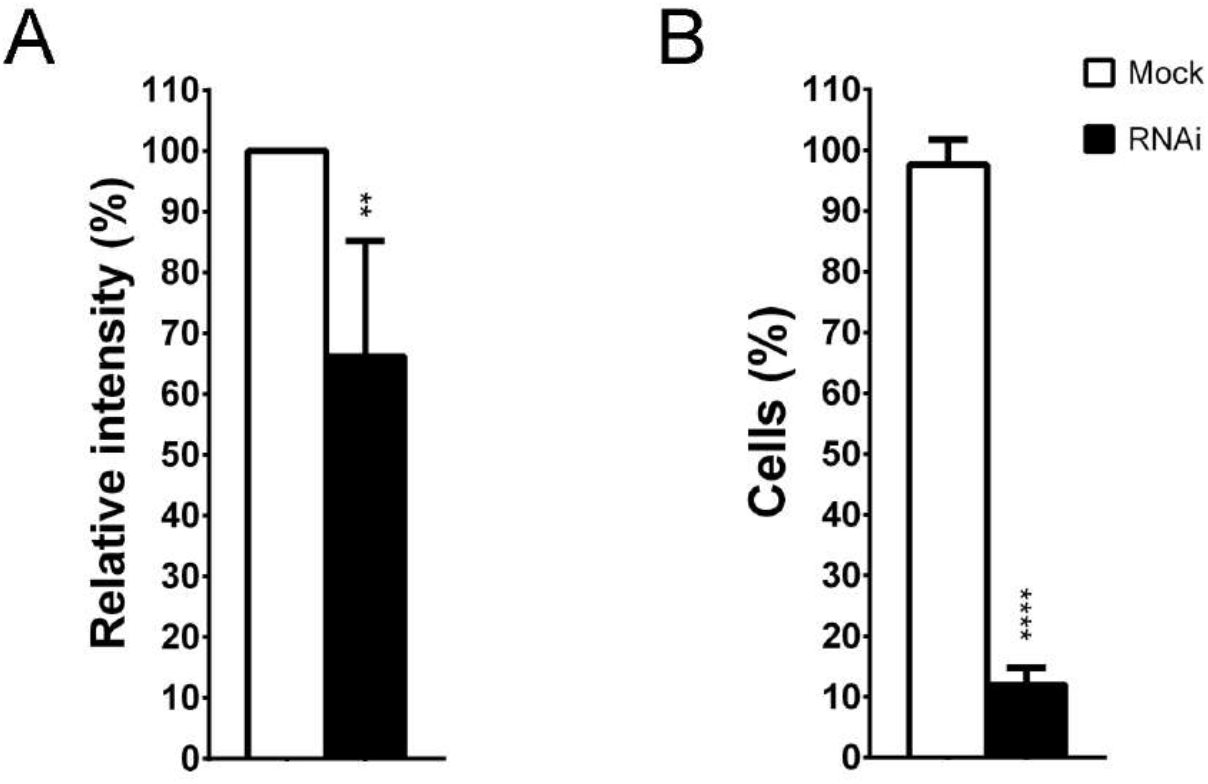
Efficiency of RNAi mediated depletion of DOM-A protein in Drosophila S2 cells. After RNAi treatments with specific DOM-A siRNAs, the decrease of DOM-A transcripts was measured by RT-PCR (A) and IF (B) and compared to control samples. Fluorescence intensity was assessed using the ImageJ software. Anti-DOM-A signal intensity clearly decreases after DOM-A depletion compared to the mock. The results are based on three experiments; the sq RT-PCR product band of mock was 66,14 ± 19 SD. Fluorescence intensity was assessed using the ImageJ software.

**Figure S6.**
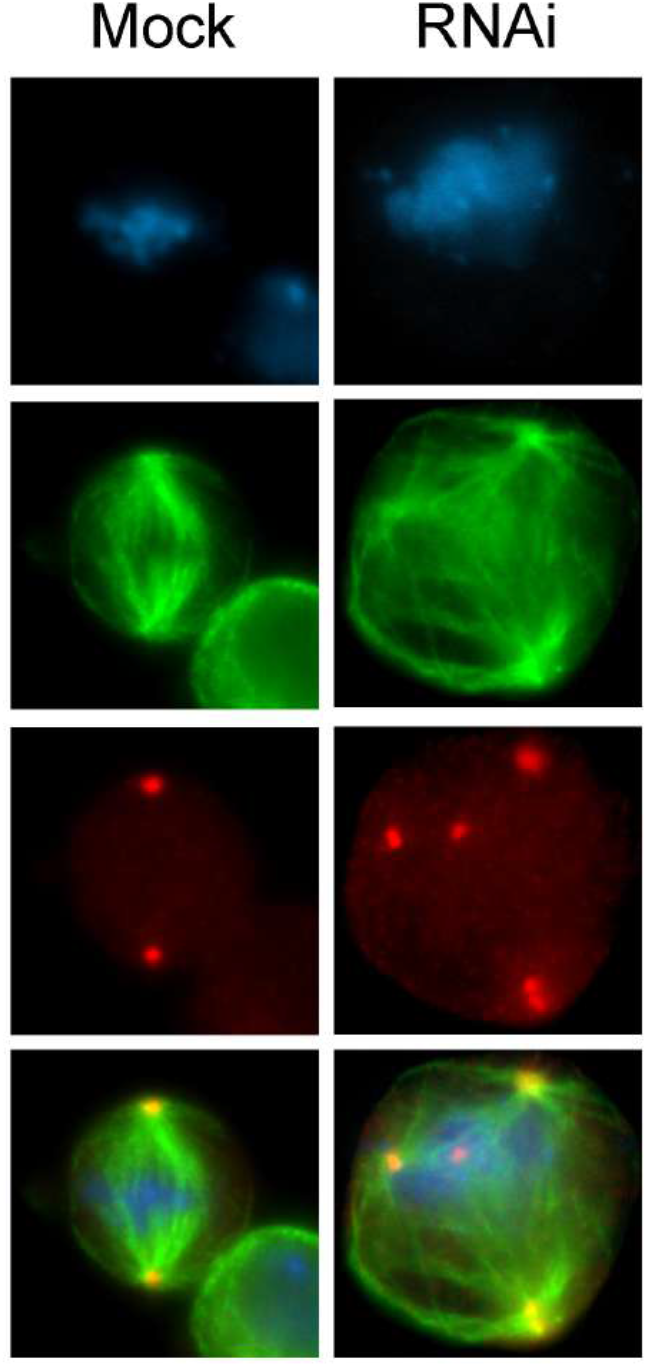
Anti-Spd2 staining of DOM-A depleted and control cells. S2 cells were stained with DAPI (blue), anti-α-Tubulin (green) and anti-Spd2 (red). In DOM-A proficient cells (mock), the anti-Spd2 staining is found on the two centrosomes at metaphase. In DOM-A depleted cells (RNAi), multiple Spd2 signals were found at multiple centrosomes which nucleate microtubules of multipolar spindles.

## SUPPLEMENTAL TABLES

**Table S1.** List of Primary antibodies

| Epitope | Reacts | Host | MW (kDa) | WB | IF | Description | Supplier | Catalog Number |
| --- | --- | --- | --- | --- | --- | --- | --- | --- |
| Alix | Human | Ms | 96 |  | 1:100 | Alix Antibody (3A9) | SantaCruz | sc-53538 |
| Anillin | Human | Rb | 130 |  | 1:500 | Anillin Antibody | Bethyl | A301-406A |
| AuroraB | Human | Rb | 39 | 1:2000 | 1:1000 | Anti-Aurora B antibody | Abcam | ab3254 |
| AuroraB | Human | Ms | 39 | 1:1000 | 1:100 | AIM-1 | BD Transduction Laboratories™ | 611083 |
| CEP55 | Human | Ms | 55 | 1:1000 | 1:100 | Anticorpo CEP55 (B-8) | SantaCruz | sc-374051 |
| Cit-K | Human | Ms | 240 | 1:2000 | 1:250 | Purified Mouse Anti-CRICK | BD Transduction Laboratories™ | 611376 |
| CREST | Human | Hu |  |  | 1:100 | Human anti-centromere/kinetochore derived from human CREST patient serum | Antibodies Incorporated | 15-234 |
| Domino A | Drosophila | Rt | 350 | 1:1000 | 1:100 |  | ML. Ruhf |  |
| H3S10phospho | Human | Rb | 18 | 1:500 | 1:100 |  | A. Losada |  |
| INCENP | Human | Rb | 120 | 1:500 | 1:100 | INCENP (P240) Antibody | Cell Signaling | 2807 |
| ISWI | Human | Rb | 150 | 1 µg/ml | 1 µg/ml |  | T. Hirano |  |
| MKLP1 | Human | Ms | 110 |  | 1:100 | Anti-MKLP1 Antibody | Abcam | ab23956 |
| MKLP2 | Human | Rb | 100 | 1:1000 | 1:250 |  | Thomas Mayer |  |
| PLK1 | Human | Ms | 70 | 1:2000 | 1:200 | Monoclonal Anti-PLK1 (STPK13) produced in mouse, clone 2G12 | Sigma-Aldrich | WH0005347M1 |
| Spastin | Human | Ms | 54/67 |  | 1:100 | Spastin Antibody (Sp 3G11/1) | SantaCruz | sc-53443 |
| Spd2 | Drosophila | Rb | 125 |  | 1:3500 |  | M.Gatti |  |
| SRCAP | Human | Rb | 350 |  | 1:100 | SRCAP (T-15) | SantaCruz | sc-133312 |
| SRCAP | Human | Rb | 350 | 1:1000 | 1:200 | C-terminal SRCAP peptide (SSAPPSLAGPAVSHRGRKAKT) | Kerafast (Chrivia) | ESL103 |
| αTubulin | Drosophila/Human | Ms | 50 | 1:20000 | 1:500 | Monoclonal DM1A | Sigma-Aldrich | T9026 |
| αTubulin | Drosophila/Human | Rb | 50 |  | 1:2000 | Anti-alpha Tubulin antibody - Microtubule Marker | Abcam | ab18251 |
| βActin | Drosophila/Human | Ms | 42 | 1:10000 |  | Beta-Actin Antibody - mouse monoclonal antibody | Abgent | AM102b |

**Table S2.** List of Secondary antibodies

| React | Conjugation | Host | Description | Dilution | Supplier | Catalog Number |
| --- | --- | --- | --- | --- | --- | --- |
| Human | FITC | Donkey | Donkey anti-human IgG (H+L) Cross-Adsorbed Secondary Antibody, Alexa Fluor 488 | 1:200 | Jackson ImmunoResearch | 709-545-149 |
| Mouse | 488 | Chicken | Chicken anti-Mouse IgG (H+L) Cross-Adsorbed Secondary Antibody, Alexa Fluor 488 | 1:200 | Invitrogen | A21200 |
| Mouse | 555 | Goat | Goat anti-mouse IgG Secondary Antibody Alexa Fluor 555 | 1:200 | Invitrogen | A21422 |
| Mouse | HRP | Goat | goat anti-rabbit IgG-HRP | 1:1000 | Santa Cruz | sc-2005 |
| Rabbit | 488 | Chicken | Chicken anti-Rabbit IgG (H+L) Cross-Adsorbed Secondary Antibody, Alexa Fluor 488 | 1:200 | Invitrogen | A21441 |
| Rabbit | 555 | Goat | F(ab')<sub>2</sub>-Goat anti-Rabbit IgG (H+L) Cross-Adsorbed Secondary Antibody, Alexa Fluor 555 | 1:200 | Invitrogen | A21430 |
| Rabbit | FITC | Goat | Anti-Rabbit IgG (whole molecule)–FITC antibody produced in goat | 1:200 | Sigma-Aldrich | F0382 |
| Rabbit | HRP | Goat | Goat anti rabbit IgG (H&L) conjugate HRP (1 mg) | 1:5000 | ImmunoReagents | GTXRB-003-DHRPX |
| Rat | FITC | Goat | Goat anti-rat IgG-FITC Secondary Antibody | 1:200 | Santa Cruz | sc-2011 |
| Rat | 550 | Donkey | Rat IgG (H+L) cross adsorbed antibody, DyLight 550 | 1:200 | Bethyl | A110-337D3 |

